## supplementary tables and figures for "*Treponema pallidum* subsp. *pallidum* with an Artificially Impaired TprK Antigenic Variation System is Attenuated in the Rabbit Model of Syphilis": File S1 (plasmid).docx

**Insert of the p*DC*arms-47p-*kan*^R^ vector (pUC57 vector backbone)**

**LEGEND**

In aqua = regions upstream and downstream of homology arms	(**NOT** **cloned** into vector)

In yellow= homology arms	(**cloned** into vector)

In green= *tp0574* promoter. In bold are -10, -35 and Ribosomal Binding Site	(**cloned i**nto vector)

In gray = kanamycin resistance gene	(**cloned** into vector)

```
Sense Primers are underlined
Antisense primers are double underlined


TGAAACACAGGCGCTCCGTTCCGTCTGTGCGCCGTGTGCGATACAGTGAGCCTTGATTCTGCGTTTGAAA
GCAGGCACAATGCGTCCCGTGCAGCGTATCATATCTTGGAATGTGAATGGAATTCGTGCCATAGAGCGGA
AAGATTTTCTCAGCTGGCTCGCGCGTGAGGCGCCTGATGTTCTCTGTTTGCAGGAGATTAAAGCGCATGA
GTCGCAGCTGAGTGCTGCGCTTCGTGCTCCGGTCTGGAGTGCTGGGGCGGGGGGTACGTACTATACCTAT
TTTCACAGTGCGCAGCGTCCTGGATACAGTGGCACGGCGCTGTTCAGTAAGCGCGCGCCAGATGCGGTGC
GTTTCTTCGGGGTTCCGGCTTTTGACTGCGAGGGGCGGATGCTTGCGGCACGCTTTGGCGAGCTGACGGT
GGTAAGCGCGTATTTTCCGAATGCGCAGGAAGGGGGCAAGCGGCTCGCGTATAAGCTTGATTTTTGCGCA
GCGTTTCGTGCGTTCTGTGATGAAGAGCGTACGGCCGGGCAGCACGTGATCTTGTGTGGTGACTACAACA
TAGCGCATAAGGAAATCGACCTGGCACATCCTCAGGAAAATGAGGGGAATCCTGGATTCCTGCCTCAGGA
GCGTGCATGGATGGATACATTTACGGAGGCAGGCTATGCGGATAGCTTCCGAGCCTTCTGCACAGAAGGG
CAGCAGTACACGTGGTGGAGCTACCGTGCCCGTGCACGCGCGCGTAACATTGGATGGCGCATCGATTACC
AGTGTGTGGACCAAGCCTTTTTAGCGCGCGTGACCTCTTCGCAGATACTGTCCGAGGTGACAGGATCGGA
TCACTGCCCAGTGTGTTTGACGTACGCGGACTAATCCGTTTCCGGGGTGAGCGGCACGTCCGCGCAAACT
AAGACGTACCCGCGCGCACAGGCAGCGTCAGAGGTGGTAGCGAACGTCCACACCCGCGGCTATGAACTGT
GCGGTGCGCGTGTTGGTCTGCTGTCTATCTTCTTCAATAATCTTTTCGCATGACCGGGGTACGCCGCTGT
ACGTGGCGCTTACCCCCAAGGACCAGTGCTCTGTCAGTTGAAAATAGCACCCCGCTGCCGCCTTGAGCAC
AAGACCGTAGTAGGTAGACGTGTAGTAATGCTGATAATTGAAGCCAGCCCCTACCGTCAGTGGCAAGCGG
ATGCGCCAGAAGGCAACCGTGTACCCGGCAGTGAGGGCAACGGGAATTGCAAGGTAATAGTACGGAGTAG
TGGGACTGTACGTATTGTTTGGATAGCTGCAATGGTACTGCACACTTGCGTCAATCCCGAGCGACAGGCC
GCGGCACACAAAGTGTTCAAACCCTAACGCCGCACTGAACGCGGGGTAGATGTACTTGTGCCCGTTGGTT
TGCGCGTTGGCATTACGGTCGTCCCCGCGCCCGCTGTTACACCAATCCACTTGAAAGAGGGGCACCGCGC
CCATGGCCGAAAGGCGTATAGTACTCCGGCCCGCCGCAGTAGTGTCCCACGGGTCCGCGTGCACCGGGTG
TGCAGCTCCCCACACTCCCGCGCATATTCCCAGCACCGGGCCGACCGCCCACCACTTCAATTGTTTCATA
CCCCGCTCCAACGCCGATCCTCTTACGCGTCTCGTCGAGGACCTACTCCATTCTACCCCCCCCCCACGGC
TGTTTGTCGAACCCTTTTTAAAGGGTTCGTTCTCGCGCGCTGGGCAGCACGCGCGTGAGGCGCCTATGCC
ATCGGGAGCTAGCGGATCCTCCCAAAAAGAGGAAGGACGCGCCTGTGTGTGCTCTGCATAAG
ACGTTGACAATCCCTGTGGGGCGTGCCTATACTCAGGCCCTCTATACGGAGGTGTAATC ATGAGCCATATTCAACGGGAGACGTCTTGCTCGAGGCCGCGATTAAATTCCAACCTGGATGCTGATTTAT
ATGGGTATAGATGGGCTCGCGATAATGTCGGGCAATCAGGTGCGACAATCTATCGATTGTATGGGAAGCC
CGATGCGCCAGAGTTGTTTCTGAAACATGGCAAAGGTAGCGTTGCCAATGATGTTACAGATGAGATGGTC
AGACTAAACTGGCTGACGGCATTTATGCCTCTTCCGACCATCAAGCATTTTATCCGTACTCCTGATGATG
CATGGTTACTCACCACTGCGATCCCCGGGAAAACAGCATTCCAGGTATTAGAAGAATATCCTGATTCAGG
TGAAAATATTGTTGATGCGCTGGCAGCGTTCCTGCGCCGGTTGCATTCGATTCCTGTTTGTAATTGTCCT
TTTAACAGCGATCGCGTATTTCGTCTCACTCAGGCGCAATCACGAATGAATAACGGTTTGGTTGATGCGA
GTGATTTTGATGACGAGCGTAATGGCTGGCCTGTTGAACAAGTCTGGAAAGAAATGCATAAGCTTTTGCC
ATTCTCACCGGATTCAGTCGTCACTCATGGTGATTTCTCACTTGATAACCTTATTTTTGACGAGGGGAAA
TTAATAGGTTGTATTGATGTTGGACGAGTCGGAATCGCAGACCGATACCAGGATCTTGCCATCCTATGGA
ACTGCCTCGGTGAATTTTCACCTTCATTACAGAAACGGTTTTTTTATAAATATGGCATTGATAATCCTGA
TATGAATAAATTGCAGTTTCATTTGATGCTCGATGAGTTTTTCTGA
TTACCATGTCACTTTCATTCCGCAGACAAGGGTGCCGAAGTGGCGTTCGGACCAGATGCTCTCGGCAATGCCCATGTAAGGAGCGTCAGCAAGCACGCCCTGTTCCCACTGGGCGCTGAGCTCCACCTTCTCGAAGGGACTGAACGTCAGTCCCACCTGGTACTGGAGCGCGTGCTCACGCAGGAGCGCATCCCCGCTCTGGTTGTGGTTGAAGCGGTTTGTTGCCCATAGCACGGATGTGTGTGGTGCAAGCCAGGCGTGGGAACCGAGGGGGATGCGATAGCTGCACCACGCTTTGCTCAAGATAGGCGTGTTGATGACGCCGTCCGAATTACTTCCCTTGTACTGCGCACCTCCGTTATTTATGTAAAAGATGTAGGTGAGGGGGATGTACACGCGTGCTTCGACGCCGGCGTTCAGGCCGGTGAGCAGGTGGGTGTAGGGGTCACCGCTTTTGGTTTCGAGTTTGAGGAATCCGGCAAAATCAAAGTGATCTGCTTGATTCTTAAAGAAAACACGTTCTCCAAAGATATTAGTGCCTGCGGTGGCAAAGTATATGCCAGAAGAGAGCCACCTCCACTGCATACGCAGGAGCGCGTCTATGTTGAGCGTTTTGGTGTACGTGAATTGCAACCAGGAAATAAAGAGTGCTGTCCGCATCCGCGCGTCAGGATTCTTCTGCTGCTCTATGAGCCTCATGTTTATTTCTAGGCCTTTGCCATTCAGATGTCTTAGGTAACCACCGCTTGACGCTAAGCCGACGATGGTGCGCTGCGCCAAGAGCGTTACTAAGCCAGCTTTCTCCAGAAGACTGCCTTTGCTCAGTTCTTGCACAAAGAATGAAACAACATCTTGTTGCGGGGAGGTCAGCAGCGTGTCCCCAAGACTAGTGATGACTTCATGGACCCTCTGTGAGGGAGCCATTCTAGCGTAGAATTGTGCGTTACTCTGGTGTTGGTTACCGG
CGTCGAGGGCGAAGGAGAAGCGGAAGCCGGCGCCTGGTTCGAGGGTGAGTCGGCCTCCTACTCCCCACAG
GAGTGCTGTTTTGTTTTCGTTCTTGGAGTCTTCGGTACCCTTAACGTAGTTCTGGTCCAGTGTGGCATTC
CCTGCCAGCTCCAACGTAAGCAGCCGCTGACGGTCGACGCCATAGGAAAGCGTTGCATCGGCCCCGAAGC
CATACTTGCTGTGCGTGGTGTCAGTACTATCCCAGGCACCATTGGAAAGGAAGGAGAGGAAACCGATGTC
CACATCTACTCCGCTGTTTCCCACATTGTGGGCCTGGTAGCCGAGTTTTGCCCCGGAGCCGGAGAAACCA
GGGGCATAGCGAGTGTCCTTTTCTGAATAGGCACGGGTGACAAAGGGTTTCCACAGCTGGGCAAAGTTAA

Green sequence primers target the tp47 promoter.
SENSE 	AGCGGATCCTCCCAAAAAGA
ANTISENSE 	GATTACACCTCCGTATAGAG

Blue seqeucne primers target T. pallidum DNA regions flanking the DCs homology arms.
SENSE		CACAGAAGGGCAGCAGTACA
ANTISENSE	CTTCTCCTTCGCCCTCGAC
```

Gray seqeunce primers: target the *kan*^R^ gene.

```
SENSE		GAGCCATATTCAACGGGAGA
ANTISENSE	ATTCCGACTCGTCCAACATC
```
