## supplementary tables and figures for "*Treponema pallidum* subsp. *pallidum* with an Artificially Impaired TprK Antigenic Variation System is Attenuated in the Rabbit Model of Syphilis": Table S1 (inoculum).docx

Table S1. TprK variability profile in the SS14 and SS14-DSKO strains used for rabbit inoculation.

| TprK variable (V) Region | Amino acid sequence* | Wild-type SS14 strain Percent sequence | SS14-DC^KO^ strain  Percent sequence |
| --- | --- | --- | --- |
| V1 | **GIASETGGAGALKH** | **96.06195** | **81.30797** |
| V1 | GIASDGGAIKH | 1.986697 | 0.355974 |
| V1 | GIASDGGAGALKH | 0.964227 | 0 |
| V1 | GIASDGGALKH | 0.705832 | 0 |
| V1 | GIASETGGAQPLKH | 0.281289 | 0.85311 |
| V1 | GIASEKNGGAGALKH | 0 | 1.265003 |
| V1 | GIAYENGGAQPLKH | 0 | 1.619613 |
| V1 | GIASEKNGGAQPLKH | 0 | 14.59834 |
| V2 | **WEGKDSKGVVQAGANHSK** | **85.1821** | **40.83901** |
| V2 | WEGKSNTGAPAAGANHSK | 7.328483 | 0 |
| V2 | WEGKPNGNVPAGANHSK | 2.010713 | 0 |
| V2 | **WEGKDSQGKAPAGANHSK** | 1.914901 | **25.66107** |
| V2 | WEGKPNGNVQAGANHSK | 1.513741 | 0.379553 |
| V2 | **WEGKSNTGVVQAGANHSK** | 1.042542 | **29.81366** |
| V2 | WEGKPNGNVPAGVTPSK | 0.606363 | 0 |
| V2 | WEGKDSKGVVQAGATHSK | 0.40116 | 0 |
| V2 | WEGKSNTGAAPAGANHSK | 0 | 0.871244 |
| V2 | WEGKSNTGAPAAGVTHSK | 0 | 0.913211 |
| V2 | WEGKSNTGAPAGANHSK | 0 | 1.522243 |
| V3 | **TLSGDYATARAGADDILWD** | **88.95463** | **65.03622** |
| V3 | TLSGGYATARAGADDILWD | 5.402198 | 0.476627 |
| V3 | TLSGDYATAQPPANILWD | 1.499912 | 0.510309 |
| V3 | TLSGDYARAGADDILWD | 1.43129 | 0.77715 |
| V3 | TLSSGYAQAAGADDILWD | 1.345246 | 1.253777 |
| V3 | TLSGDYATARAGAAVPAAADDILWD | 0.722128 | 0 |
| V3 | TLSGDYATAPANNDILWD | 0.644596 | 0 |
| V3 | **TLSGDYAQAAGAGADDILWD** | 0 | 28.20839 |
| V3 | TLSGDYATARARPPAPANNAILWD | 0 | 0.233595 |
| V3 | TLSGDYAVAPADDILWD | 0 | 0.276496 |
| V3 | TLSSGYAQAAAAAVNNDILWD | 0 | 2.335733 |
| V3 | TLSSGYAQAARAGADDILWD | 0 | 0.891698 |
| V4 | **TDVGRKKDGAQGTV** | **96.91317** | **96.16694** |
| V4 | TDVGHKKENAANVNGTV | 1.534279 | 0.516221 |
| V4 | TDVGHKKNGANGDI | 0.664887 | 1.094878 |
| V4 | TDVGRKKDGANGDI | 0.348438 | 1.395811 |
| V4 | TDVGHKKDGAQGTV | 0.273043 | 0.293197 |
| V4 | TDVGRKKDGAQGSV | 0.266185 | 0 |
| V4 | TDVGHKKENAANVKGTV | 0 | 0.266477 |
| V4 | TDVGRKKGGAQGTV | 0 | 0.266477 |
| V5 | **QASNVFQGVFLTDTTPMLQHDC** | **53.176** | 1.716312 |
| V5 | **QASNVFQGVFLTTPMQKDDC** | **26.09685** | 0.443286 |
| V5 | **QASNVFQGVFLTTPMLQHDC** | **12.33243** | **68.85708** |
| V5 | QASNVFQGVFLTDTTPMQKDDC | 1.709207 | 0 |
| V5 | KASNVFEGVFLARNIAMLQHDC | 1.540043 | 0 |
| V5 | **KASNVFKDVFLTNAMDMQTHDC** | 1.511896 | **12.80529** |
| V5 | QASNVFQGVFLTTPMQKHDC | 1.281905 | 0 |
| V5 | QASNVFQGVFLTNNMLQHDC | 1.007427 | 1.74132 |
| V5 | QASNVFKDVFLTNAMDMQTHDC | 0.764118 | 0.483447 |
| V5 | QASNVFQGVFLTNAMDMQTHDC | 0.580125 | 0.749419 |
| V5 | **KASNVFKDVFLTDTTPMQTHDC** | 0 | **10.32818** |
| V5 | KASNVFKDVFLTNNMQTHDC | 0 | 0.766091 |
| V5 | KASNVFKDVFLTTPMQTHDC | 0 | 0.861565 |
| V5 | KASNVFQGVFLTDTTPMQTHDC | 0 | 0.804732 |
| V5 | QASNVCQGVFLTTPMLQHDC | 0 | 0.443286 |
| V6 | **PVHWKALAPAQPPARVDIY** | **49.2956** | 0 |
| V6 | **PVHWKALARAQPPARVDIY** | **27.46295** | 0 |
| V6 | PVHWKALARAVPPARVDIY | 3.354095 | 0 |
| V6 | PVHYKVLKAHAQPPARVDIY | 2.127645 | 0 |
| V6 | PVHWNAFTQARALPGAPVPAIY | 1.808006 | 2.825492 |
| V6 | PVHWNAFTQAQPPARVDIY | 1.746048 | 0 |
| V6 | PVHWKALAPAQPPANIY | 1.673526 | 0 |
| V6 | PVHWKALARARPPARVDIY | 1.570739 | 3.589618 |
| V6 | PVYYFAARAQPPARVDIY | 1.419366 | 0 |
| V6 | PVHYTVLTGPQAPAAANIY | 1.288407 | 0 |
| V6 | PVYYFAARAQLPAVAPANDIY | 1.117334 | 0 |
| V6 | PVHYTVLTGPQPPARVDIY | 0.869503 | 0 |
| V6 | PVYYFAAPAQPPARVDIY | 0.814588 | 0 |
| V6 | PVHWKALAPALPPARVDIY | 0.715322 | 0 |
| V6 | PVHWKALAPAQPPARVDIN | 0.684344 | 0 |
| V6 | PVHYKVLKAHARAPADIY | 0.639279 | 0 |
| V6 | PVHWKALAPARPPARVDIY | 0.553385 | 0 |
| V6 | PVHAQARAQPPANIY | 0.440034 | 0 |
| V6 | PVHYKVLKAHARAPADIH | 0.440034 | 0.88187 |
| V6 | PVHWKALARALPPARVDIY | 0.432991 | 0 |
| V6 | PVHWKALAPAQPPAAANIY | 0.416098 | 0 |
| V6 | PVHWKALPAAAAAVNNDIY | 0.405533 | 0 |
| V6 | PVHWKALAPAQPPARGDIY | 0.378076 | 0 |
| V6 | PVHYTVLTGPQAPAAANIN | 0.347097 | 0.463066 |
| V6 | PVHALARARPPVPAIY | 0 | 0.305874 |
| V6 | PVHAQARPPVPAIY | 0 | 0.305874 |
| V6 | PVHWKALAPAQPPAIY | 0 | 0.745391 |
| V6 | PVHWKALAPARPPVPAIY | 0 | 0.373684 |
| V6 | PVHWKALAQARPPVPAIY | 0 | 1.023162 |
| V6 | PVHWKALARAGAAVPAIY | 0 | 0.763267 |
| V6 | PVHWKALARALPGAPVPAIY | 0 | 1.963216 |
| V6 | PVHWKALARALPPAIY | 0 | 0.285162 |
| V6 | PVHWKALARALPPVPAIY | 0 | 0.446048 |
| V6 | PVHWKALARAQPPAIY | 0 | 0.498818 |
| V6 | PVHWKALARAQPPVPAIY | 0 | 0.593872 |
| V6 | PVHWKALARARPLPPAIY | 0 | 0.249409 |
| V6 | PVHWKALARARPPAPAIY | 0 | 1.249882 |
| V6 | **PVHWKALARARPPVPAIY** | 0 | **75.82605** |
| V6 | PVHWKALARAVPLPPAIY | 0 | 0.516695 |
| V6 | PVHWKALARAVPPVPAIY | 0 | 0.350135 |
| V6 | PVHWKAQARALAAGVPAIY | 0 | 0.30871 |
| V6 | PVHWKAQARARPPVPAIY | 0 | 0.403764 |
| V6 | PVHWNAFTQAQPPAIY | 0 | 0.287998 |
| V6 | PVHWNAFTQARPPVPAIY | 0 | 0.828241 |
| V6 | PVHYKVLKAHARAPVPAIY | 0 | 0.463066 |
| V6 | PVHYKVLKAHARPPVPAIY | 0 | 1.827855 |
| V6 | PVYYFAARAQLPAIY | 0 | 0.445189 |
| V6 | PVYYFAARAQLPAPVPAIY | 0 | 0.630484 |
| V6 | PVYYFAARAQLPAVAPAIY | 0 | 0.466761 |
| V6 | PVYYFAARARPPVPAIY | 0 | 0.647501 |
| V6 | PVYYFAARPPVPAIY | 0 | 0.433845 |
| V7 | **YGGTNKKNDAAPAAPATKWKAEYCGYY** | **56.19608** | **41.41053** |
| V7 | **YGGTNKKNDAAPATKWKAEYCGYY** | **38.03072** | 3.803587 |
| V7 | YGGTNKQAAPAAPATKWKAEYCGYY | 1.446126 | 0 |
| V7 | YGGTNKKAAAAAPAPATKWKAEYCGYY | 1.133211 | 0 |
| V7 | YGGTNKKATPPAAPAAPATKWKAEYCGYY | 0.681696 | 0 |
| V7 | YGGTNKKAAPAAPATKWKAEYCGYY | 0.647046 | 0 |
| V7 | YGGTNKKNDAAPAAPATKWKAGYCGYY | 0.571029 | 0.559721 |
| V7 | YGGTNKKNDAAPAAPTKWKAEYCGYY | 0.480513 | 0.484263 |
| V7 | YGGTNKKNDAAPAAPATKWSKEYCGYY | 0.431364 | 1.265345 |
| V7 | YGGTNKKNDAAPAALTKWKAEYCGYY | 0.382215 | 0 |
| V7 | YGGTNKKNDAAPAAPGTKWKAEYCGYY | 0 | 0.638955 |
| V7 | YGGTNKKATPPAAPAAPTKWKAEYCGYY | 0 | 0.949644 |
| V7 | YGGTNKKNDAAPAAPGTKWSKEYCGYY | 0 | 1.664056 |
| V7 | YGGTNKKNDAATKWKAEYCGYY | 0 | 0.500604 |
| V7 | **YGGTNKQAATKWKAEYCGYY** | 0 | **46.71209** |
| V7 | YGGTNKQAATKWSKEYCGYY | 0 | 2.011204 |

* Highly represented sequences (>10% of the total) are in bold.
