## supplementary tables and figures for "*Treponema pallidum* subsp. *pallidum* with an Artificially Impaired TprK Antigenic Variation System is Attenuated in the Rabbit Model of Syphilis": Table S3 inoculum variants lower than 0.25%.docx

| **Table S3. Inoculum variants detected in rabbits infected with the SS14-DC^KO^ strain at <0.25% reads.** | | | | |  |
| --- | --- | --- | --- | --- | --- |
| V region | Nucleotide Sequence | Amino acid sequence | Coordinates on SS14 genome (NC 021508.1/CP004011.1)^1^ | Sample^2^ (Percent reads) | Percent reads in inoculum |
| V2 | TGGGAGGGCAAAGACAGTCAGGGCAAGGCCCCAGCAGGAGTAACCCCCAGCAAG | WEGKDSQGKAPAGVTPSK | 152027-151989 | KO R503 B03 Day30 (6.4) | 0.107 |
| V3 | ACACTGTCTAGCGGCTATGCCCAAGCAGCCGGAGCCGGAGCCGACGACATCTTATGGGAT | TLSSGYAQAAGAGADDILWD | 150743-150701 | KO R504 B04 Day37 (0.3) | 0.171 |
| V3 | ACGCTTTCTGGCGACTATGCCCAAGCAGCCCGAGCCGGAGCCGACGACATCTTATGGGAT | TLSGDYAQAARAGADDILWD | 150743-150701 | KO R504 B04 Day37 (0.3) | 0.068 |
| V5 | AAGGCCAGCAATGTGTTTAAAGACGTCTTTCTCACCGATACCACACCCATGCTGCAGCACGACTGT | KASNVFKDVFLTDTTPMLQHDC | 152277-152213 and  148544-148483 | KO R503 B02 Day23 (0.2) | 0.224 |
| V5 | AAGGCCAGCAATGTATTTCAGGGAGTATTTCTCAACATGGCCATGACCGCACACGACTGT | KASNVFQGVFLNMAMTAHDC | 151684-151627 | KO R501 B02 Day23 (4.9)  KO R501 B03 Day30 (23.3) | 0.114 |
| V5 | AAGGCCAGCAATGTATTTCAGGGTGTCTTTCTCACCACACCCATGCTGCAGCACGACTGT | KASNVFQGVFLTTPMLQHDC | 149794-149742 | KO R501 B02 Day23 (0.2) | 0.080 |
| V5 | AAGGCCAGCAATGTGTTTAAAGACGTCTTACTCACCGATACCACACCCATGCAGACGCACGACTGT | KASNVFKDVLLTDTTPMQTHDC | 152277-152213 | KO R501 B05 Day41 (0.4)  KO R503 B02 Day23 (0.4)  KO R503 B03 Day30 (0.5)  KO R504 B02 Day23 (0.4)  KO R507 B03 Day30 (0.5) | 0.072 |
| V5 | AAGGCCAGCAATGTGTTTAAAGACGTCTTACTCACCAATGCCATGGACATGCAGACGCACGACTGT | KASNVFKDVLLTNAMDMQTHDC | 152277-152213 | KO R501 B05 Day41 (0.3)  KO R503 B04 Day37 (0.5)  KO R504 B04 Day37 (0.5)  KO R504 B07 Day58 (0.7)  KO R504 B08 Day65 (0.8)  KO R507 B02 Day23 (0.3)  KO R507 B04 Day37 (0.3) | 0.063 |
| V6 | CCCGTCCATTGGAACGCCTTCACCCAAGCCCTACCCCCAGCCATCTAC | PVHWNAFTQALPPAIY | 148727-148691 | KO R503 B06 Day44 (0.4) | 0.024 |
| V7 | TACGGCGGTACGAACAAGAAAAACGATGCTGCTCCTGGTACGAAGTGGAGCAAGGAATATTGTGGGTATTAC | YGGTNKKNDAAPGTKWSKEYCGYY | 159529-159574 and 149577-149543 | KO R507 B05 Day41 (18) | 0.128 |
| V7 | TACGGCGGTACGACCAAGAAAAACGATGCTGCTCCTGCTGCTCCTGCTACGAAGTGGAAGGCAGAATATTGTGGGTATTAC | YGGTTKKNDAAPAAPATKWKAEYCGYY | 149013-148971 | KO R503 B02 Day23 (0.2)  KO R503 B04 Day37 (0.2) | 0.110 |
| ^1^Coordinates of the DC that contains the sequence detected in this V region.  ^2^KO = rabbit infected with the DC^KO^ strain; R = rabbit#; B = biopsy#; Day# = day post-inoculation; (%) = percentage of sequence compared to all sequences detected in the sample for a given V region at this time point. | | | | |  |
