## Supplementary figures and images for "*Treponema pallidum* subsp. *pallidum* with an Artificially Impaired TprK Antigenic Variation System is Attenuated in the Rabbit Model of Syphilis"

### Fig.S1_V1_heatmap_min5_025.pdf

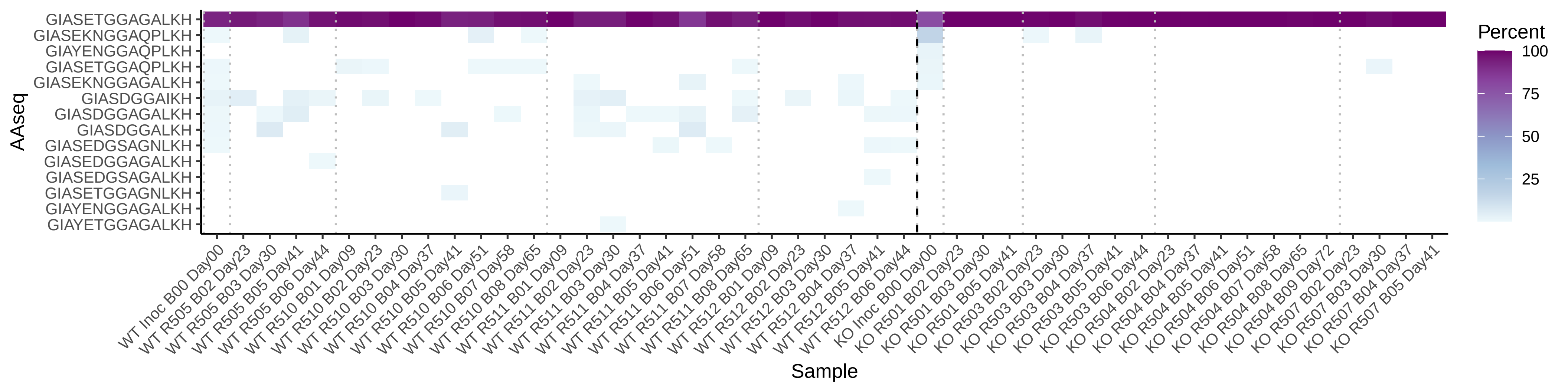

### Fig.S2_V2_heatmap_min5_025.pdf

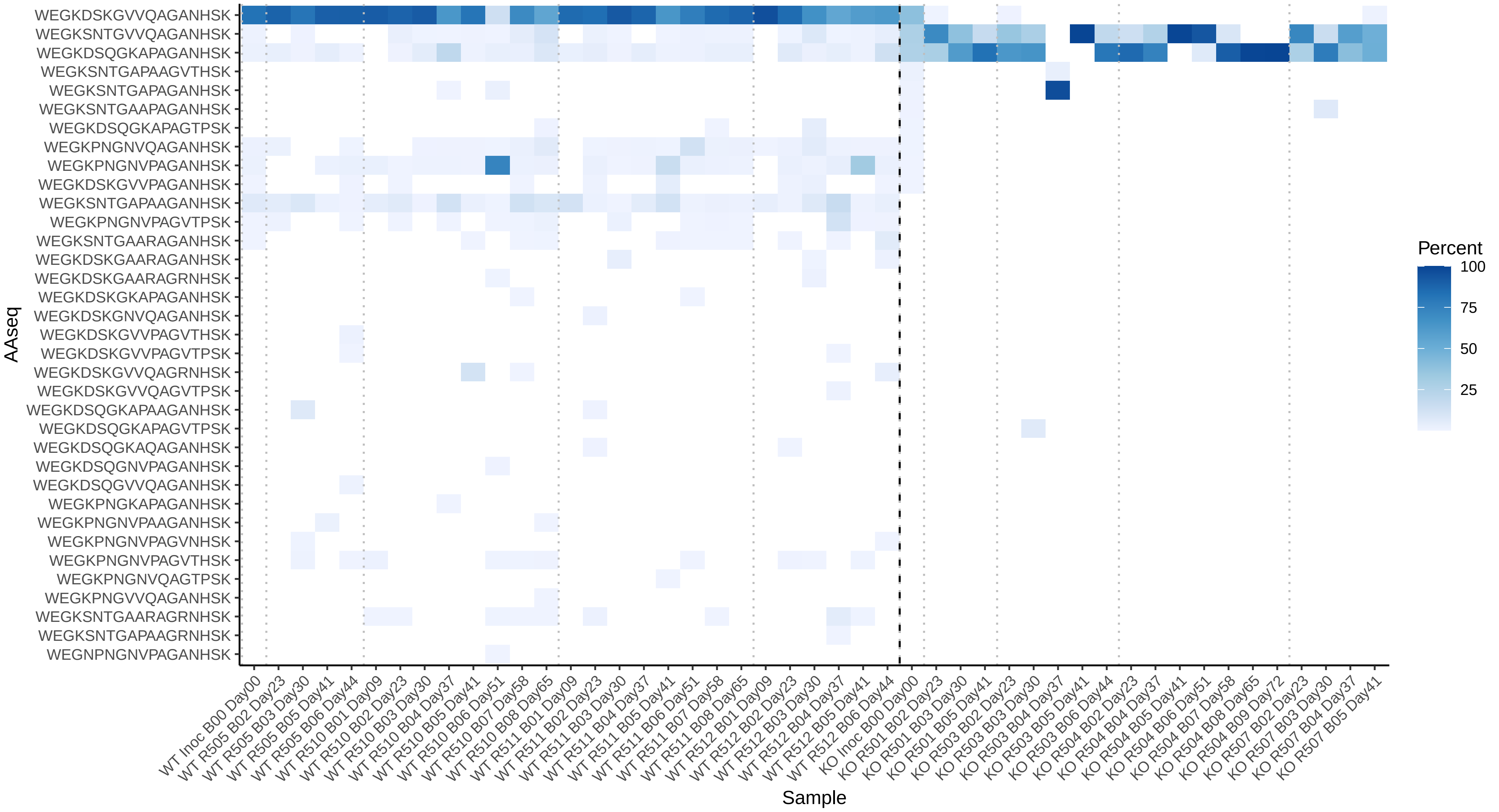

### Fig.S3_V3_heatmap_min5_025.pdf

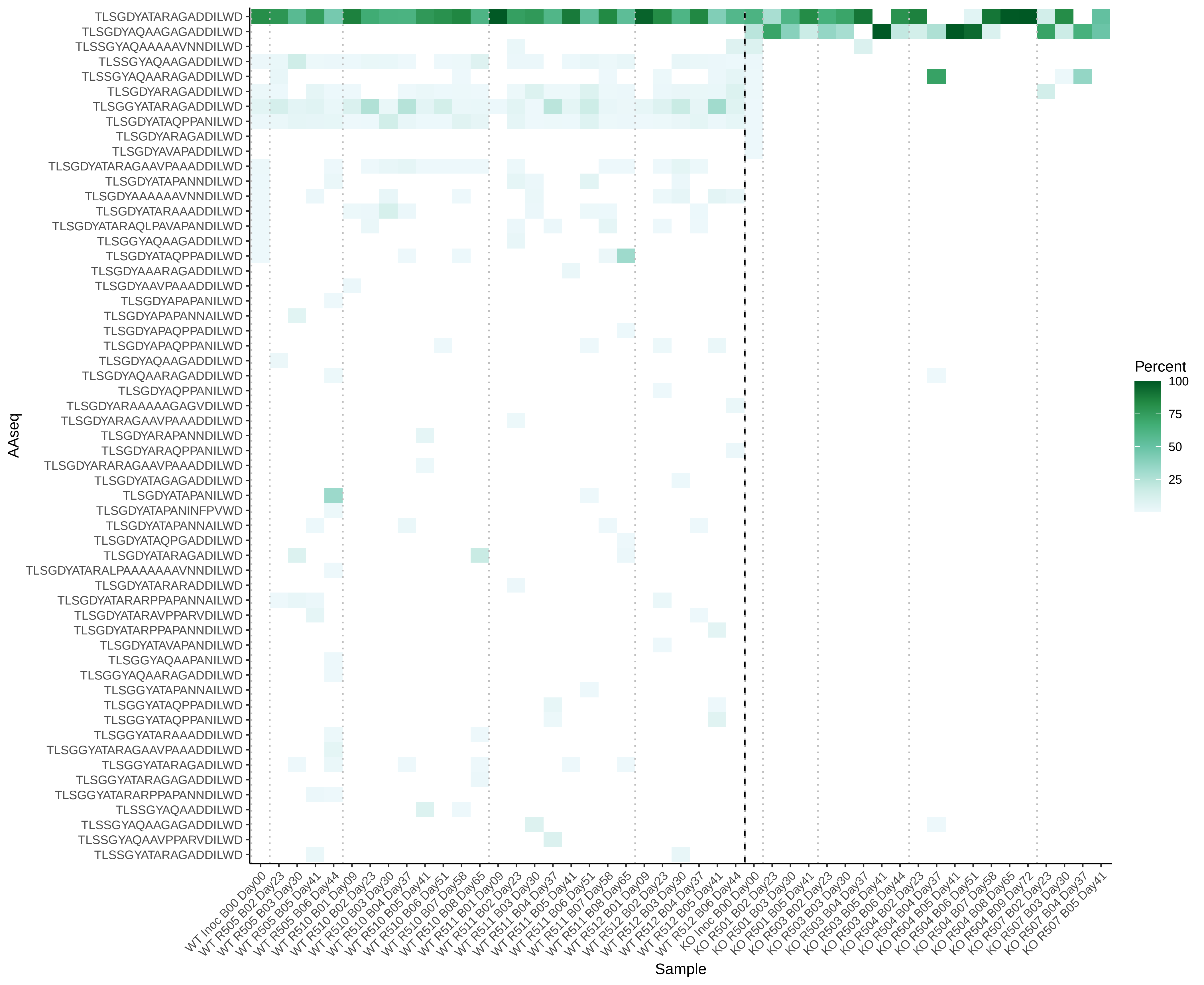

### Fig.S4_V4_heatmap_min5_025.pdf

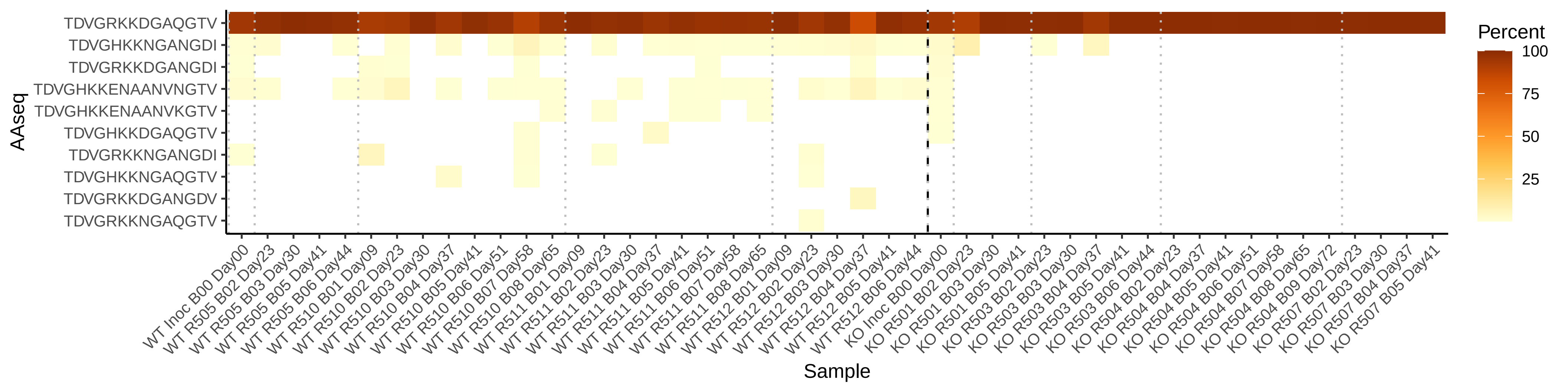

### Fig.S5_V5_heatmap_min5_025.pdf

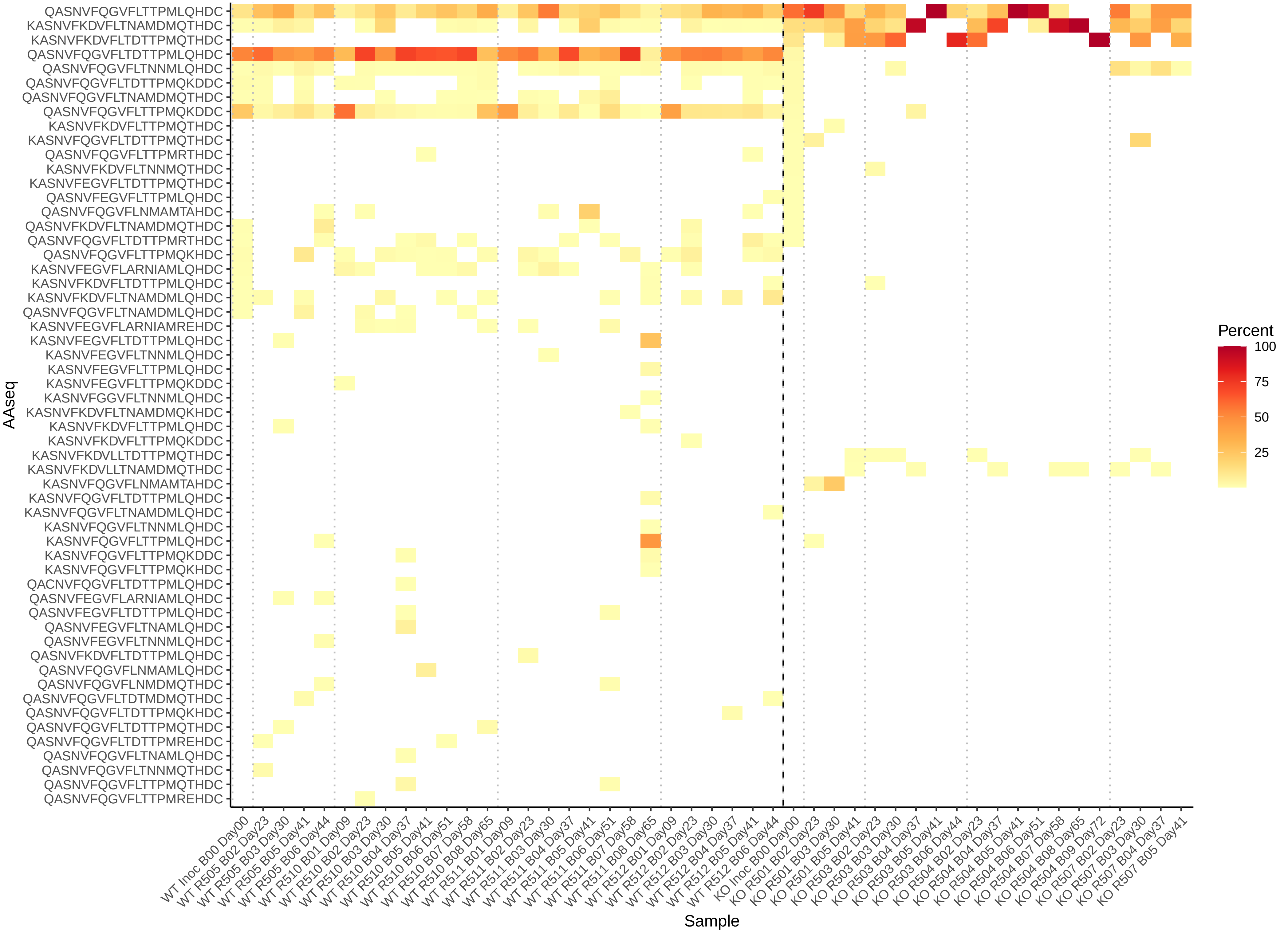

### Fig.S6_V7_heatmap_min5_025.pdf

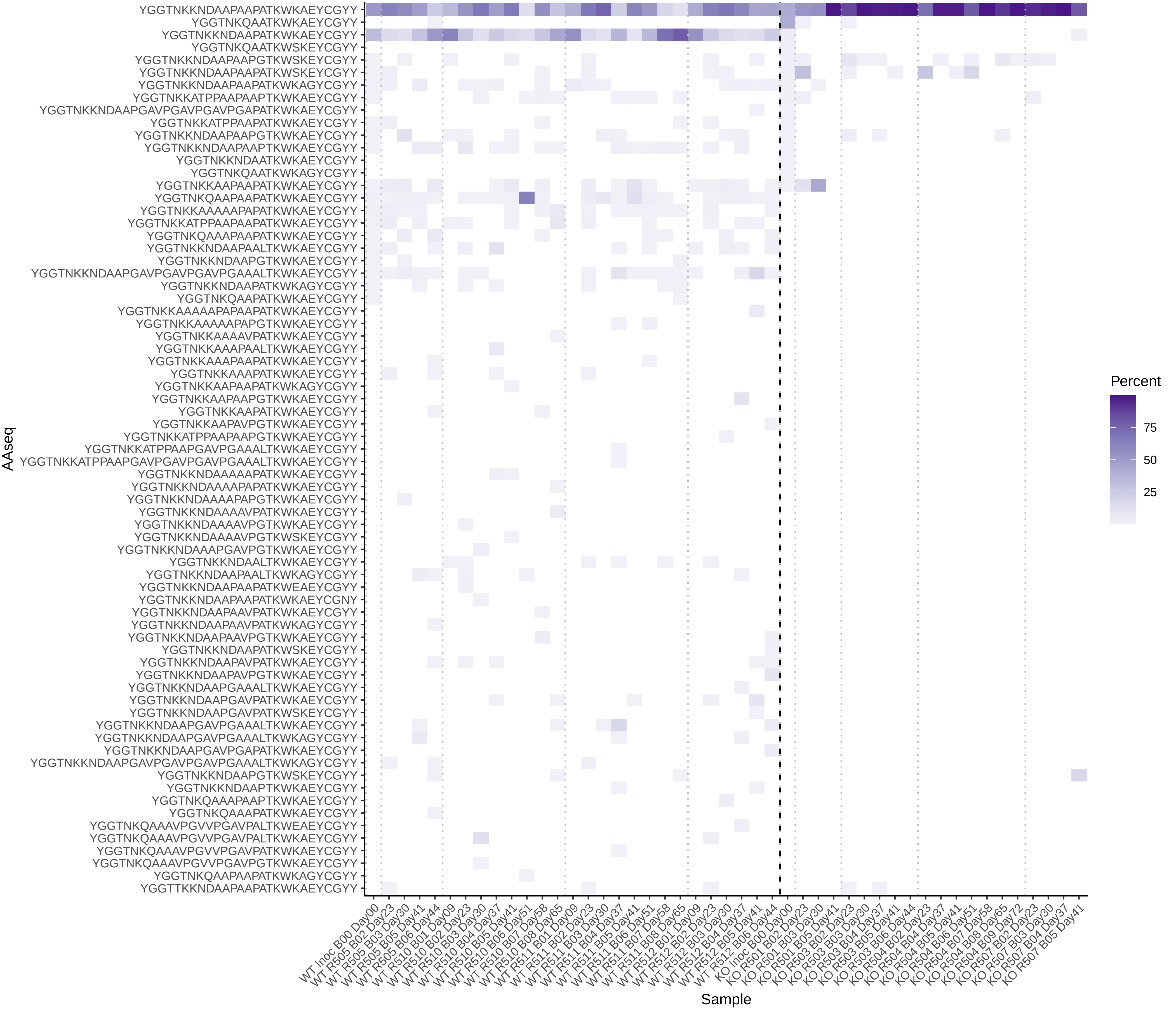
